# Synthesis, Physiochemical and Biological evaluation of Inclusion Complex of Benzyl Isothiocyanate encapsulated in cyclodextrins for triple negative breast cancer

**DOI:** 10.1101/2021.02.14.430873

**Authors:** Shivani Uppal, Rajendra Kumar, Khushwinder Kaur, Shweta Sareen, Alka Bhatia, S.K. Mehta

**Author notes:** These authors contributed equally to this work (Shivani Uppal, Rajendra Kumar). S. K. Mehta, Department of Chemistry and Centre of Advanced Studies in Chemistry, Panjab University, Chandigarh-160 014, India.

## Abstract

Benzyl isothiocyanate (BITC), an organic dietary compound, is allied with a major role in the potential chemopreventive effects. BITC has acknowledged rising attention as a therapeutic compound to be used in medicine because of its high potency and characteristic biopharmaceutical properties, like high permeability with marginal aqueous solubility. The highly volatile and hydrophobic nature brought a need to provide a suitable delivery-matrix to BITC to exploit its pharmacological potential to the fullest. It has been successfully incorporated in β-CD and HP-β-CD using acoustic forces and thoroughly characterized using UV-vis spectroscopy, FTIR, DSC, TEM, and SAXS. The complexation helped in masking the acute odour, achieving a controlled release of BITC, and made its use viable by prolonging the retention time and thereby sustaining the biological effects. Different models like Higuchi, first-order kinetic decay, Korsmeyer-Peppas model were applied, suggesting a diffusion-controlled mechanism of release. Also, the bioaccessibility and stability of BITC in an *in vitro* digestion model was evaluated. The main objective of the present work was to systemically study the credibility of BITC-CD complexes in well-established tumor mimicking 2D cell culture models and produce a conclusive report on its chemotherapeutic activity. The *in vitro* anti-cancer activity of BITC and the formed sonochemical complexes was confirmed by MTT assay and further evaluated using apoptosis assay and production of ROS like moieties. Cell cycle analysis was done to evaluate the growth inhibitory mechanism of BITC. Strikingly, BITC and its complexes showcased ROS generation and lysosome-mediated cell death. Effect on cell migration was assessed using wound healing assay. The results promptly suggest the functional efficacy of the CDs in releasing BITC and attest the ability of the complexes to provide alternate to otherwise remedially sparse triple-negative breast cancer.

## INTRODUCTION

Cancer, scientifically known as “malignant neoplasm,” is a broad group of diseases involving unregulated cell growth. The reason behind the origin of this fatal disorder has been a compelling question for decades leading to various theories like humoral theory, lymph theory, chronic irritation theory, etc., only to discover the oncogenes as the leading cause of cancer (1, 2). Breast cancer is the most common in women and the second most common cause of cancer death in women in the U.S (3). The BRCA1 gene accounts for 5to 10% of all inherited breast cancers, and inherited mutations of the BRCA1 gene account for about 40–45% of hereditary cancers (4). Clinically, the most useful markers in breast cancer are the estrogen, progesterone, and HER-2/Neu receptors that are used to predict response to hormone therapy (5). “Triple-negative breast cancer (TBNC)” is a special category of breast cancer accounting for about 10-20%, which does not express any of these receptors (6). Patients with TBNC have higher mortality rates compared to other subtypes of cancer, probably due to a lack of validated molecular targets and a lesser understanding of this cancer. Conventional chemotherapy has been tried for a long to treat breast cancer, and its partial success is the leading cause that researchers are still exploring alternatives to replace the stereotyped completely or to supplement it to achieve the pinnacle (7, 8).

The medical field has grown by leaps and bounds in the last few decades, bringing new hope to the patients of breast cancer (9, 10). The numerous treatments like chemotherapy, lumpectomy, radiation therapy, adjuvant therapy, etc., available for the cure of early detected breast cancer are highly expensive and are followed by a series of side effects. So, there has been a growing interest in finding more economical and human-friendly cures and prevention treatments of cancer. Plant and herbs derived chemicals (10) are described to play an essential role in prevention and are now being tried for therapeutic capabilities. Among various natural antitumor agents, isothiocyanates, especially, Benzyl isothiocyanate (BITC) purified from cruciferous vegetables, have shown to be capable of bearing the burden of tumor treatment (9). BITC is an important member of the isothiocyanate family, a biologically active dietary phytochemical found in the Eastern Hemisphere *Alliariapetiolata*, seeds of the *Salvadorapersica*, and *Carica papaya*. It is a yellow liquid with a characteristic watercress-like odor. It is investigated significantly to explore the chemopreventive and antimicrobial properties. It can not only inhibit chemically induced cancer but has also shown promising results against oncogenic-driven tumor formation and human tumor xenografts in rodent cancer models (9). Besides, BITC is also found to be clinically useful in the management of breast cancer (11). However, viable administration of these potent plant derivatives to achieve an optimum adaptive cellular stress to deliver the clinically defined setpoints has mostly remained unanswered. The biologically imperative properties of BITC are hindered due to its volatility, high photosensitivity, and fragility. One of the major challenges with using BITC is the lack of solubility in aqueous-based body tissue fluids, which hinders its penetration into cellular levels and results in low bioavailability and poor therapeutic efficiency. BITC with proven cytotoxic effects requires a suitable carrier system that can deliver it slowly at tumor sites. Reports are scarce (12-14) that have investigated the plausible improvement of the biopharmaceutical potential of BITC. It, therefore, calls for the adoption of more effectual and novel delivery strategies for enhancing the bioavailability and bio-accessibility of BITC.

Cyclodextrins (CDs) are glucopyranose units with a hydrophobic interior and a hydrophilic exterior. They can potentially augment the biological availability of BITC due to their pivotal benefits like ease of formulation development, high encapsulation efficiency, long-term stability, and ease of industrial scalability (15). CDs can simultaneously serve the function of a carrier; improve the organoleptic properties by curbing acute odor and solubility enhancer, making them an impeccable choice for nutraceutical formulations. The controlled release of BITC from the CDs can effectively suppress the proliferation rate of cancer cells by extending the retention time of BITC (12). Thus, the aptness of CDs as the delivery wagon for BITC was first examined using physicochemical characterization followed by biochemical and cellular functions in well-established tumor mimicking models. In the present approach, an attempt has been made to evaluate the potential of CDs as a carrier to cargo potent BITC inside the breast cancer cells. The in-depth analysis of *in vitro* assessment of six cellular processes (cytotoxicity, cellular uptake, cell death mechanism, cell cycle arrest, ROS production, measurement, and migration) on the MDA-MB-231 has been undertaken. Also, owing to the high mortality rate of Triple-negative Breast cancer, MDA-MB-231cells were purposefully used to survey the ability of a pristine BITC or encapsulated form of BITC to penetrate and kill these cells known to have resistance towards many other chemotherapeutic agents.

## MATERIALS AND METHODS

Benzyl Isothiocyanate, β-CD, hp-β-CD, hexane, Dulbecco’s eagle medium (DMEM), Phosphate Buffered Saline, were procured from Sigma-Aldrich, purity > 99%), while Ethanol (absolute) was obtained from Changshu Yangyuan Chemical (China). Millipore water with conductance less than 3 µS cm-1 was used for all the preparations. MDA-MB-231cells were obtained from NCCS, Pune, India. Fetal bovine serum (FCS), Antibiotic/antimycotic cocktail, Trypsin-Ethylenediaminetetraacetic acid (EDTA) solution, L-Glutamine, MTT, Propidium Iodide (PI) were purchased from Himedia, India. Annexin-PI apoptosis kit was procured from Thermo Fisher Scientific, USA.

### Preparation of Inclusion Complexes

The inclusion complex of BITC with β-CD (β-CD BITC) and with HP-β-CD (HP-β-CD BITC) was prepared by using a probe sonication technique. The method has been reported in our previous work (12). Briefly, 0.044 mol of BITC was added dropwise to an equimolar solution of β-CD (or HP-β-CD) with a minimal solvent mixture (ethanol:water::2:8). The mixture was heated until60 °C under continuous magnetic stirring. The solution was sonicated for 900 s (3 sets of 300 s each) at 180 W using Hielscher UP200Stultrasonic devices. The end products were obtained by lyophilization. The amount of BITC entrapped in the ICs was determined spectrophotometrically.

### Spectroscopic and thermal characterizations

UV-visible (UV) absorption spectra were recorded using the JASCO V-530 spectrophotometer (4-21, Sennin-cho 2-chome, Hachioji, Tokyo 193-0835, Japan model). The spectral range (190-450nm) was covered with a precision of ± 0.2 nm, using quartz cells having a path length of 1 cm. FTIR spectra were recorded with thermally controlled diode laser in the spectral region of 4000-500 cm^-1^ using Thermo scientific, Nicolet iS50 FT-IR. The thermal properties of the formed complexes were investigated using differential scanning calorimetry (DSC) Q20 (M/s TA Instruments, Detroit, USA). Each prepared sample (5–8 mg) was heated in a crimped tin pan for DSC. All samples were heated from room temperature to 350°C at 10 °C/ min under a nitrogen flow of 40 ml/min. The mass loss and heat flow in the sample were recorded as a function of temperature with reference to an empty pan. Reproducibility was checked by running the sample in triplicate.

### Size and Morphological Analysis

The morphology and size of the formed complexes were analyzed using Hitachi H-7500 Transmission electron microscopic (TEM) using a 300-mesh copper grid. The grid was kept in an inverted state and a drop of the prepared formulation was applied to the grid for 10s. Excess of the formulation was removed by absorbing on a filter paper, and the grid was analyzed using Hitachi (H-7500) (Japan) 120 kV equipped with CCD camera with a resolution of 0.36 nm (point to point) and 40–120 kV operating voltage.

Small Angle X-ray Scattering (SAXS) measurements were recorded on the SAXSess mc^2^ instrument (Anton Paar, Austria) using a line collimation system at 20 °C and exposure times of 30 minutes. The powdered sample was introduced between two scotch tape sheets, fixed in a powder cell before being placed into the evacuation chamber. The scattered intensities were recorded by the Eiger detector at 307 mm distance from the sample. The scattering intensity I(q) was measured as a function of q =4π/λ sin θ, with 2θ as the scattering angle, λ = 0.154 nm as the wavelength of the used radiation and range of scattering vector as 0.06-10 nm^-1^. All the data were corrected for the background scattering from the scotch tape and the slit smearing effects by a de-smearing procedure using SAXSAnalysis software employing the Lake method. The estimations of the size of the concerned systems were done utilizing the Pair distance distribution function p(r) using the Generalized Indirect Fourier Transformation (GIFT) program, whereas the electron density profiles were obtained using the DECON program from the PCG software package^5,6^(Anton Paar, Austria). The electron density profiles have been computed from the Pair distance distribution function p(r) under the assumption of spherical symmetry.

### Release Studies and Kinetics

*In vitro* release studies were conducted by employing the dialysis bag method at two different pH, i.e., 5.5 and 7.4 in phosphate buffer saline (PBS). Initially, the dialysis bag was equilibrated in dissolution media for 3 h. The formulations containing bioactive equivalent to 20 ppm of BITC were put in the dialysis bag (MWCO: 4kDa, Himedia, India). The temperature for release studies was kept constant, i.e., 37 ± 0.1°C with the help of a thermostat. The dialysis bag was dipped in the vessel containing 50 mL of dissolution media under gentle stirring at 100 rpm. At fixed intervals of times, withdraw 2 ml of sample from the dialysis jacket followed by replenishment with an equal volume of temperature-equilibrated PBS. The concentration of the collected samples was determined spectrophotometrically at λ_max_254 nm using the calibration curve formed under similar conditions. Various kinetic models were applied to study the kinetic behavior and release mechanisms of BITC from the inclusion complex. The results obtained from release studies were fitted into different kinetic models such as zero-order kinetics, first-order kinetics, Higuchi kinetics, Korsmeyer and Pappas (KP) and Hixson-Crowell model represented as Equations. (1-5) (16).

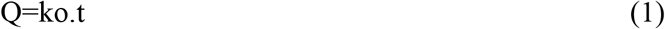

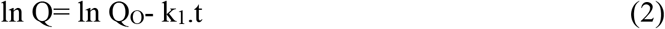

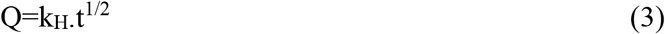

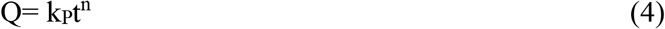

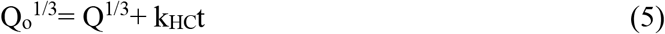

Where, Q_o_is the initial amount of BITC in the system, Q denotes the fraction of drug released up to time t, k_o_, k_1,_ k_H_, k_P_, k_HC_ are the constants of various models and n is the release exponential that describes the release kinetics.

### Bio-accessibility and stability of BITC in an *in vitro* digestion model

A simulated GIT model based on a previous study [2] was used to compare the potential gastrointestinal fate of the β-CD BITC and HP-β-CD BITC complexes. Simulated saliva fluid (SSF), simulated gastric fluid (SGF), and simulated intestinal fluid (SIF) were prepared as described in the previous study (17). Briefly, SSF imitates the biochemical environment of the mouth phase which primarily consists of a mixture of mucin and various salts. SGF mimics the acidic settings present in the stomach by the addition of NaCl and HCL in a container with 1L double-distilled Millipore water. Finally, the SIF comprises CaCl_2_ and NaCl in the molar ratio of 1:15 in the presence of a lipase. After *in vitro* digestion, 1 mL of the raw digest was centrifuged (12000 rpm) for 30 min. The supernatant with solubilized BITC was collected. BITC in the raw digest and the micelle phase was extracted with ethanol and n-hexane. The extracted BITC was then analyzed using JASCO V 530 spectrophotometer. The stability and bio-accessibility of BITC after digestion were measured according to a reported method (18).

### Cell line studies

MDA-MB-231 (an epithelial, human breast cancer) cells were obtained from NCCS Pune with a recommendation to grow in Leibovitz media. Cells were routinely cultured and harvested from the culture flasks (T-25 cm^2^; BD Falcon, USA), maintained as per ATCC guidelines. Subsequently, these cells were passaged and cryopreserved. Further, cryopreserved cells were thawed and from that point forward were grown in DMEM media supplemented with 2mM L-glutamine, 1X antibiotic/antimycotic cocktail, 10% FBS. Cells were cultured as a monolayer at 37 °C in a thermostat-controlled incubator (Thermo Scientific Midi 40) having 5% CO_2_ and 90% relative humidity and passaged once reached 80-90% confluency. Passaged cells were used for various experiments after seeding in apposite culture vessels.

### *In-vitro* cytotoxicity study

The *in vitro* cytotoxicity of free BITC, β-CD-BITC, and HP-β-CD BITC was evaluated using3-[4,5-dimethylthiazol-2-yl]-2,5-diphenyltetrazoliumbromide (MTT) assay by a previously reported standard method (19). Briefly, 5×10^3^ MDA-MB-231 cells were seeded in each well of 96 well plates (BD Falcon, USA) and were kept in an incubator overnight to allow attachment. The next day, free-floating cells were removed, and the media was replaced with the one containing different dilutions (200 to 0 ppm) of all three treatments. β-CD-BITC and HP-β-CD BITC concentrations were adjusted to contain an equivalent concentration of BITC. Cells were incubated for 48 h, and after incubation, the medium containing treatment was removed and replaced with a medium containing 10% MTT (5mg/mL) and were kept for 4 h. Finally, MTT supplemented media was removed, and formazan crystals so formed were dissolved by adding 100 µL DMSO per well. The optical density of purple color was measured by using a microplate reader (Epoch 2 Microplate Spectrophotometer by BioTek) at 570 nm. Untreated cells were used as a control to calculate the % cytotoxicity. IC_50_ values for all three treatments were calculated using GraphPad Prism Software. Cell viability was expressed as 100% untreated cells (control). All samples were performed in triplicate, and the survival rate was calculated with the following equation (6).

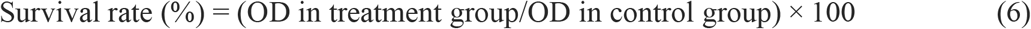

### Cellular uptake

Cellular uptake of free BITC, β-CD-BITC, and HP-β-CD BITC was evaluated using flow cytometry (BD AccuriC6, California, USA). Briefly, 5 × 10^5^ per wells were seeded in each well of a six-well culture plate (Thermo Fisher Scientific) and kept overnight. The next day, attached cells were treated with 10 ppm equivalent concentration of BITC. Cells were washed and detached using trypsin: EDTA and analyzed immediately on a Flow cytometer.

### Cell death Mechanism Study

The cell death mechanism was studied using the Annexin-V-Propidium iodide (PI) method (20). Briefly, 1×10^5^ cells were seeded in each well of 6 wells culture plate and were incubated overnight. The cells were treated with BITC, β-CD-BITC, and HP-β-CD BITC containing 2.5, 5, 10, and 20 ppm equivalent concentration of active drug moiety. Cells were treated for 48 h, and after incubation, medium containing cells were centrifuged to rescue floating cells. The attached cells were collected by trypsinization and were pooled with floating cells of respective treatment. After removing the culture medium completely using centrifugation, cells were resuspended in 100 µL 1x annexin binding buffer. Cells were stained with Annexin V-Alexafluor 488 and PI as per manufacturer protocol (Thermo Fisher, USA) for 15 minutes. After 15 min, 300 µL annexin binding buffer was added to the tubes containing stained tubes, and cells were immediately analyzed on a flow cytometer.

### Cell Cycle Analysis

DNA cell cycle analysis was done on treated with BITC, β-CD-BITC, and HP-β-CD BITC. After 48 h of treatment with 2.5, 5, 10, and 20 ppm BITC equivalent concentrations, cells were removed by trypsinization, washed, and fixed with 70% chilled ethanol. Cells were stored in 70% ethanol for few days in a refrigerator till flow cytometric analysis. For analysis, cells were suspended in PBS containing 50 µg/mL RNase and 20 µg/mL propidium iodide. Cells were stained for 15 minutes at 37 °C and were analyzed on a Flow cytometer. Doublets were removed using height vs. area plot for propidium iodide.

### Intracellular Reactive oxygen species (ROS) measurement

Production of ROS was measured using 2’-7’-Dichlorodihydrofluorescein diacetate (DCFH-DA) on a flow cytometer. Cells were seeded in a six-well plate at a density of 1×10^5^ cells per well and cultured overnight. The next day, cells were treated with 2.5, 5, 10, and 20 ppm BITC equivalent concentrations for 12 hr. After completion of incubation, cells were washed, and 10 µM 6-carboxy-2′,7′-dichlorodihydrofluorescein diacetate (DCFH-DA) containing serum-free media was added for 10 min incubation at 37 °C in a fluorescent ROS indicator (21). Once done, cells were trypsinized and washed with PBS before acquisition on a flow cytometer. Cell-associated fluorescence was measured at 533/33 nm fluorescence channel. Untreated cells were used as a control to measure the basal level of ROS, and data were presented as a histogram showing mean fluorescence intensity (MFI).

### Lysosomal tracking assay

Lysosomes are known to induce apoptosis by the discharge of diversity of hydrolases constituted by them, which are released from lysosomal lumen to cytosol in response to a variety of stimuli, including oxidative stress. Lysosomal enrichment, which can be linked to cytotoxic response, was measured by flow cytometric analysis of cells treated with BITC, β-CD-BITC, and HP-β-CD for 2.5, 5, 10 and 20 ppm BITC equivalent concentrations for 12 hr. After treatment, cells rinsed to remove media containing respective treatment and were stained with 50 nM LysoTracker Red for 30 minutes. After staining, cells were washed twice with PBS, followed by flow cytometric analysis.

Similarly, in a separate set, cells were stained with Acridine orange for microscopic analysis of lysosomes. For microscopic analysis, brightfield and Acridine-orange, red fluorescence images were captured on an inverted fluorescent microscope (Nikon, Japan). Microscopic images were analyzed and overlaid using ImageJ software.

### Wound healing scratch assay

Cellular migration accompanied by continuous proliferation is known as a hallmark of cancer cells. Scratch assay was used to monitor this behavior of MDA-MB-231 cells after treatment with free BITC, β-CD-BITC, and HP-β-CD BITC. Cells were grown until the monolayer was formed and were serum-starved overnight. The next day, cell monolayers were scratch using 200 µL volume dispensing sterile micropipette tip to create a scratch. Cells so freed were removed by gentle rinsing with sterile prewarmed PBS. After this cell was incubated with 5, 10, and 20 ppm BITC equivalent concentrations for 24 h and images were captured at 0, 6, 18, and 24 h post-treatment. Free BITC was not assayed in scratch assay because of the high toxicity of free BITC. Cells treated with media with 10 % FBS was used as a positive control for cell migration. Images were acquired by keeping in mind that the same area for the given treatment set was imaged at each time point. All the images were analyzed using T-Scratch.

### Statistical Analysis

Results are expressed as the mean ± standard deviation of at least three independent experiments. Graphical representation also includes standard deviation as vertical error bars. Further, measurements were statistically analyzed using either Student’s t-test or Mann Whitney test post normality testing. The level of significance was set at p <0.05 and the power of the test at 0.8.

## RESULTS AND DISCUSSION

### Characterization and size analysis

UV-vis absorption measurements were performed to determine the molecular encapsulation of BITC in two CDs, i.e., β-CD and HP-β-CD. These studies proved to be very suitable to explore the structural changes and formation of inclusion complexes (22). UV-visible absorption spectra of BITC showed single broadband centered at around 254 nm (23). The absorption peaks of β-CD BITC and HP-β-CD BITC were deformed, and absorbance value decreased as compared to native BITC (Fig.1 (A, B). The studies on sonochemical complexes further revealed not only simple, superposition but also a slightly shifted peak at 254 nm. Although there was nearly no influence of each CD on the peak wavelength of BITC, there were considerable changes in the absorbance at the peak wavelength. This might be taken as indirect proof of complex formation (24).

FTIR was used to determine the interaction between CDs and the guest molecule in the solid-state (25). Inclusion complexes can be testified by the modification of the peak shape, position, and intensity (14). FTIR spectra of BITC, β-CD, HP-β-CD, and the complexes were collected in 4000-500 cm^-1^ region (Fig. 1C, D). Due to the vibration of the N=C=S group, the IR spectra of BITC was characterized by the peak at 2040 cm^-1^ (26). In the spectrum of the β-CD (Fig.1C), the band at 3385 cm^-1^ represents the vibration of symmetrical and asymmetrical stretching of the OH groups, and another band at 2925 cm^-1^ is associated with the vibration of C–H stretch. The absorption band at 1645 cm^-1^ corresponds to bending of H–O–H, while the bands at 1157 and 1029 cm^-1^ are associated with the vibrations of the asymmetric stretch of the C–O–C and symmetric stretching link C–O–C, respectively (27). The FTIR spectra of HP-β-CD depicted in Fig.1C is similar to the spectra of β-CD showing prominent absorption peak at 3200-3400cm^-1^ (for O-H stretching vibrations). The presence of water in HP-β-CD resulted in the presence of a broad peak of –OH. It was responsible for the reduction in wavenumber of the peak located at 2040cm^-1^. This absorption peak was masked in the inclusion complex because the aromatic ring present in the molecule was inserted in the hydrophobic cavity of cyclodextrin during the complex formation. This confirmed the inclusion and the formation of the inclusion complex (28). On the other hand, the FTIR spectrum of β-CD BITC and HP-β-CD BITC showed modification of signals associated with BITC and CDs. By comparison, the spectra of β-CD BITC and HP-β-CD BITC was not completely congruent with β-CD and HP-β-CD, respectively. The band located at 1088, 1183, 1325, 1544, 1623cm^-1,^ and 2040 cm^-1^ had shifted and diminished. Important changes in the characteristic bands of pure substances evidenced the formation of the inclusion complexes (29).

**Figure 1:**
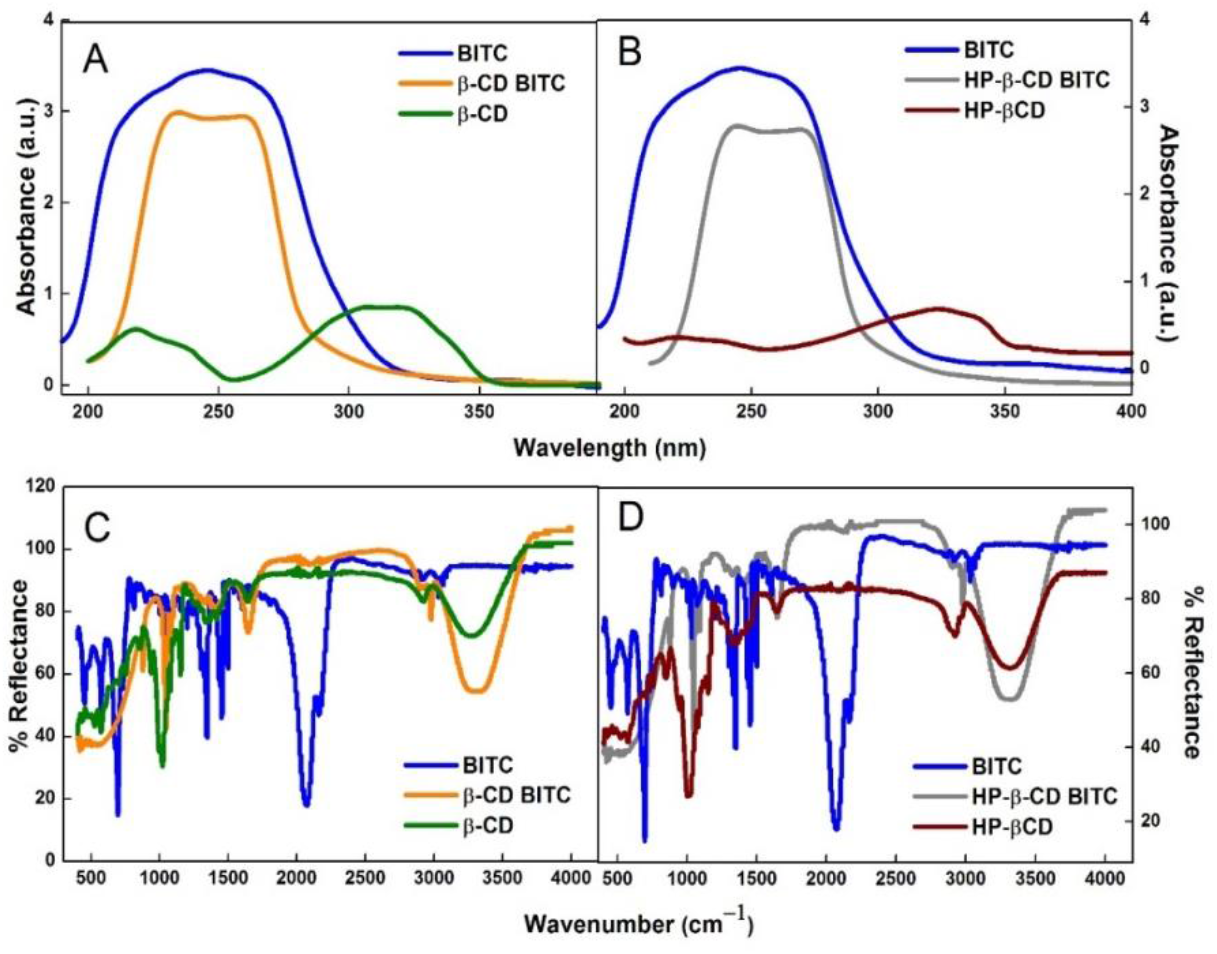
UV-Vis absorption of (A) BITC, β-CD BITC andβ-CD (B) BITC, HP-β-CD BITC and HP-β-CD; FTIR (C) BITC, β-CD BITC and β-CD (D) BITC, HP-β-CD BITC and HP-β-CD

The DSC thermograms illustrated in Fig. S1 (Supplementary Information) provide another evidence for the formation of inclusion complexes with BITC. The inclusion of the guest molecules into the host cavity leads to the changes in the physical properties of both guest and host most molecules, and therefore, the peaks corresponding to the crystal lattice, melting, boiling, and sublimation points either get shifted or disappears (29). The obtained DSC of BITC exhibits a typical sharp peak at 271°C, correspondings to the boiling point of native BITC. The thermogram of β-CD, HP-β-CD showed wide peaks around 115 °C owing to the release of water. The thermogram of the sonochemically formed complexes showed a peak shift at around 105 °C. The shift of peak to the lower temperature and intensification/deepening of the peak around 220 °C could be attributed to the change in the properties followed by complete complexation thus, confirming the interaction between BITC and CDs (30). The high thermal stability of the complexes was proved by the non-isothermal thermogravimetric analysis conducted in our previous work (29). The analysis also gave evidence of the stoichiometric ratio of host: guest to be 1:1 in the formed complexes.

The inherent morphological and size aspects of the optimal sonochemical complexes were mapped using TEM. Histograms were plotted using Image J software to give a graphical representation of the average sizes obtained (Fig. 2 (A, B). Both the complexes were found to be spherical in shape with some irregular boundaries. The images depict the particles to be well-dispersed with curtailed signs of aggregation. The average size of the complexes was evaluated to be 39 ± 2 nm and 42 ± 1.5 nm for β-CD BITC and HP-β-CD BITC, respectively.

**Figure 2:**
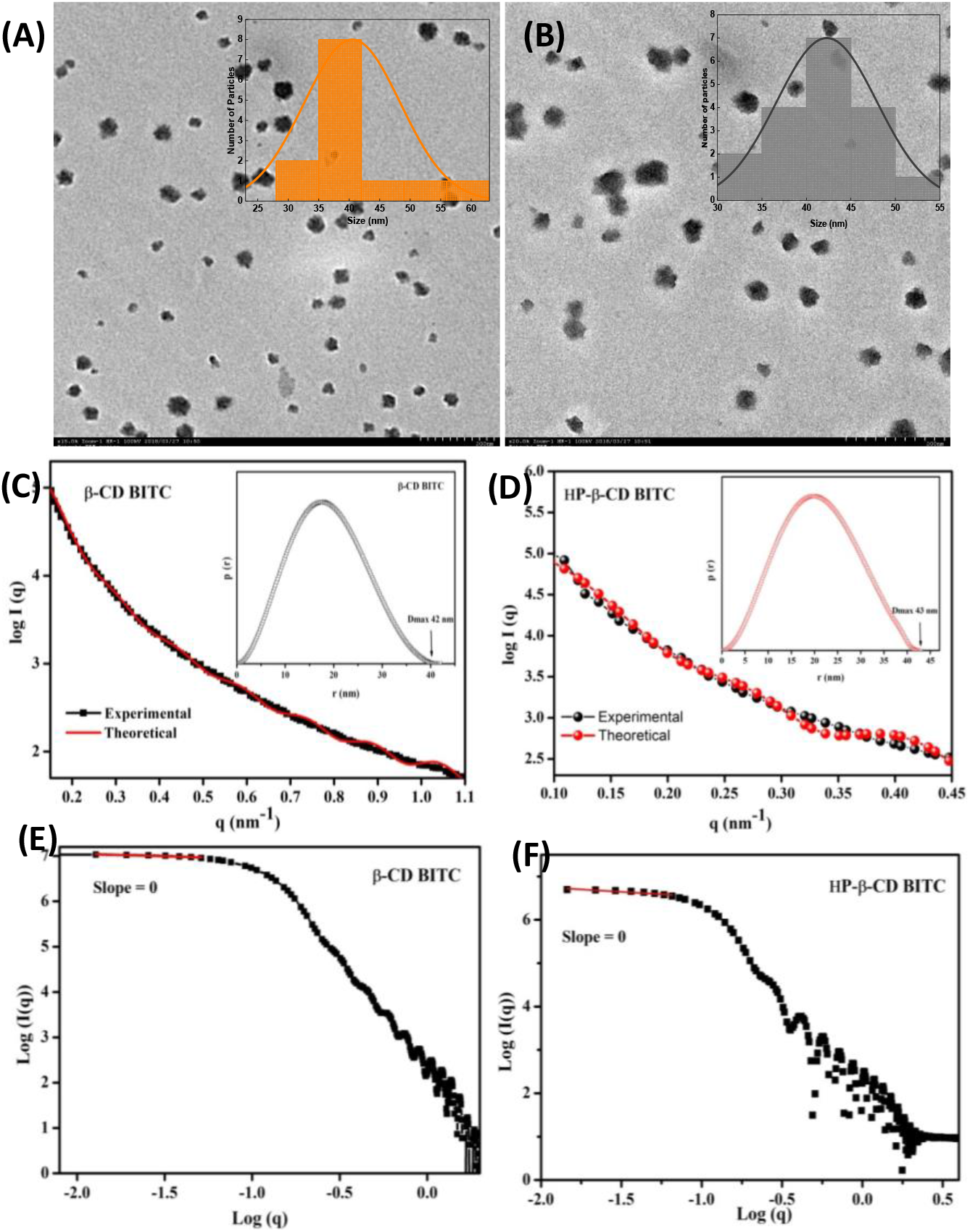
TEM Images of (A) β-CD BITC (B) HP-β-CD BITC; Scattering profiles and the PDDF distribution (Inset) of (C) β-CD BITC and (D) HP-β-CD BITC; Double logarithmic plots of the prepared complexes (E) β-CD BITC and (F) HP-β-CD BITC

To further evaluate the size and the shape of the sonochemically prepared systems, the complexes were investigated by SAXS. The corresponding scattering plots and pair-distance distribution function (PDDF) P(r) are represented in Fig. 2 (C, D). The PDDF function represents a histogram of the distances inside the particle, giving information about the maximum dimension (D_max_) as well as the shape of the particle (31). The curves exhibited a bell shape with pronounced maxima, thus leading to the fact that the particles in the system seem to be spherical with a total dimension of 42 and 43 nm for β-CD BITC and HP-β-CD BITC, respectively. Interestingly, an increase in the size of HP-β-CD complex is observed, signifying that the interactions between the primary particles are strengthened due to interaction between the hydrophobic cavity of HP-β-CD and BITC leading to the complex formation (32), which is also in consonance with the IR and TEM studies. Moreover, a double logarithm plot, i.e., log(I) *vs*. log(q) plot (Fig. 2(E, F)), confirm the shape of the scattering components that can be inferred from the slope of the curve (33). In both the cases, β-CD BITC and HP-β-CD BITC complexes, the slope of the curve was equal to zero, illustrating that the scattering components in the complexes have a spherical shape in agreement with the PDDF curves (34). The corresponding electron density plots (Fig. S2, Supporting Information) of both the complexes depicted similar profiles except for the increase in size that is exhibited by r = 21.5 nm for HP-β-CD BITC complex (also verified through the maxima of the PDDF curve) in comparison to the r = 21 nm for β-CD BITC complex.

### Release studies and kinetics

Understanding the release profile is very important for real-time investigations of any biological formulation. The *in vitro* release experiments were conducted at two pH to investigate the successful inclusion and the sustained release characteristic of BITC from the inclusion complexes. The release profile of native BITC has been reported in our earlier work (35) and is given in Fig. S3. It shows complete release in less than 4 hours. This stimulates the need for complexation. It was done to increase the solubility and provide a sustained effect to the release of BITC, which is vital in drug delivery to prolong the effect of the drug. The pH values in normal physiological environments, cell endosomes, and cells were 7.4, 5.5–6.0, and 4.0–5.0, respectively. BITC is more soluble under acidic conditions; the solubility and hence the release increases at pH 5.5. The cumulative release profiles (Fig. 3) showed a release of 54.45 and 62.45% for β-CD BITC at pH 7.2 and 5.5, respectively, while HP-β-CD BITC complex showed the release of 66.1 and 76.23% at pH 7.2 and 5.5, respectively. The release equilibrium was reached after ∼6 h for all the tested samples with a maxim drug released concentration of 10.89 mg/L (pH 7.2) and 12.49 mg/L (pH 5.5) for β-CD BITC complex. However, the maxim drug released concentration of 13.33mg/L (pH 7.2) and 15.24 mg/L (pH 5.5), respectively for HP-β-CD BITC complex. Similar attainment of equilibrium at low release percentages for BITC has been observed by Parmar *et al*. (36) (∼53% release of BITC from PLGA Nps at pH 7.4) and Kumar *et al*. (37) (51% release of BITC from α-tocopherol based oil in water NEm). In comparison to β-CD, HP-β-CD exhibited a much stronger controlled release ability for BITC. The higher cumulative release of HP-β-CD BITC complex than β-CD BITC complex can be explained by the combinatorial advantages of BITC and HP-β-CD complexation and higher solubility of HP-β-CD as compared to β-CD. This further indicates that the interactions between the BITC and HP-β-CD are relatively weaker than β-CD BITC complex when the intermolecular forces are neutralized in a buffered salt solution; as a result, higher release is observed.

**Figure 3:**
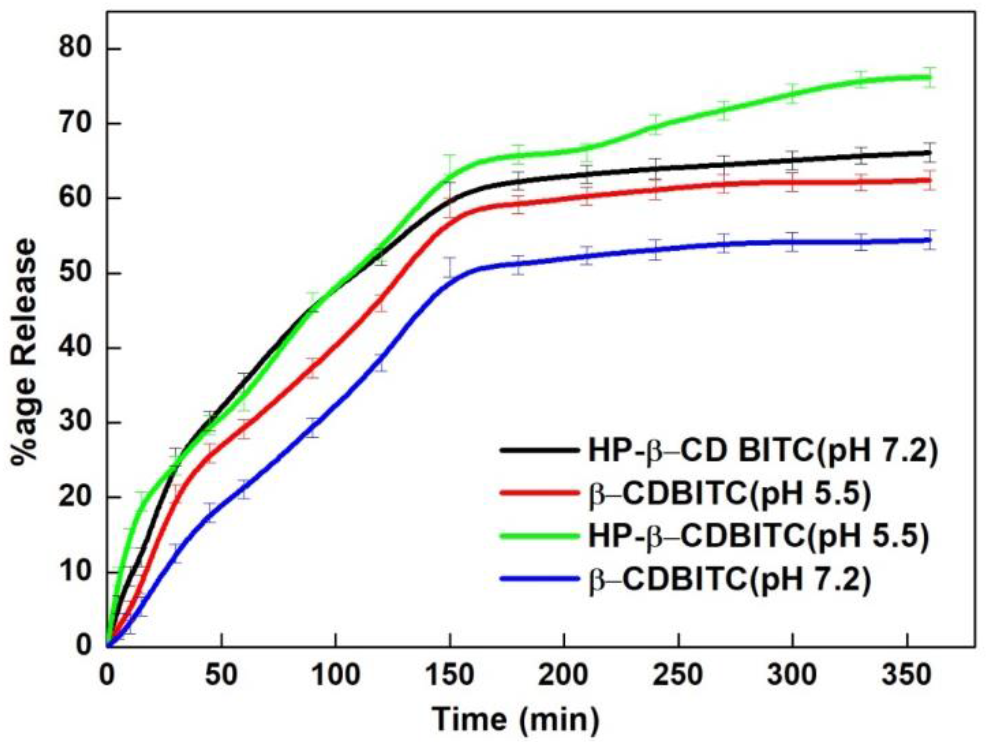
Release profiles of β-CD BITC and HP-β-CD BITC complex at pH 5.5 and 7.2

To understand the release kinetics ofβ-CD BITC and HP-β-CD BITC complexes, different kinetic models (Eqs.1-5) were applied to the drug release profiles. Table S1 (Supplementary information) lists values of the rate constants and regression correlation (R^2^) calculated using the rate equations for the release of BITC at pH 7.2 and 5.5. The mechanism of release was well explained by KP models in terms of the diffusion exponent (n) and constant (k_P_). The Fickian release was observed for both formulations.

### Bioaccessibility and stability of BITC in an *in vitro* digestion model

The application of BITC as a healing bioactive is established, but no real-time data can be obtained to demonstrate the true potential of the molecule. One of the reasons for this is the low solubility and chemical instability in the gastrointestinal tract. This, in turn, leads to low bioavailability and accessibility. So, the stability and sequential digestion in the mouth, stomach, and intestine to access the bioaccessibility of both the complexes - β-CD BITC and HP-β-CD BITC were evaluated using the method reported by Liu *et al*. (17). The analysis of the digesta collected after undergoing extreme changes in pH and ionic strength was done in the last stage of the *in vitro* model. The results documented the stability of BITC to be slightly higher in β-CD BITC (73.45 ± 1.75%) than in HP-β-CD BITC (69.96 ± 2.73%). This is due to better thermodynamic stability (prominent factor) of β-CD as compared to HP-β-CD. Further, the effective bio-accessibility was calculated, and it was found to be 61.34 ± 2.59% and 58.54 ± 2.72% for β-CD BITC and HP-β-CD BITC, respectively. The bio-accessibility values are directly dependent on the stability of BITC inside the complexes. By comparison of both complexes, the obtained results clearly show that it is expedient to encapsulate BITC in CDs. The slight difference observed may be explained on the basis of different colloidal stability of complexes under simulated digestion conditions. Similar results have been obtained by Chen *et al*. (38). Our findings indicate that the nano-complexation between CDs and BITC provides a promising strategy to improve the stability and bio-accessibility of BITC.

### Cytotoxicity evaluation

Comparative effects of pure BITC and both the complexes (β-CD BITC and HP-β-CD BITC) were investigated on MDA-MB-231 breast cancer cell lines using MTT assay (39) (Figure 4).

**Figure 4:**
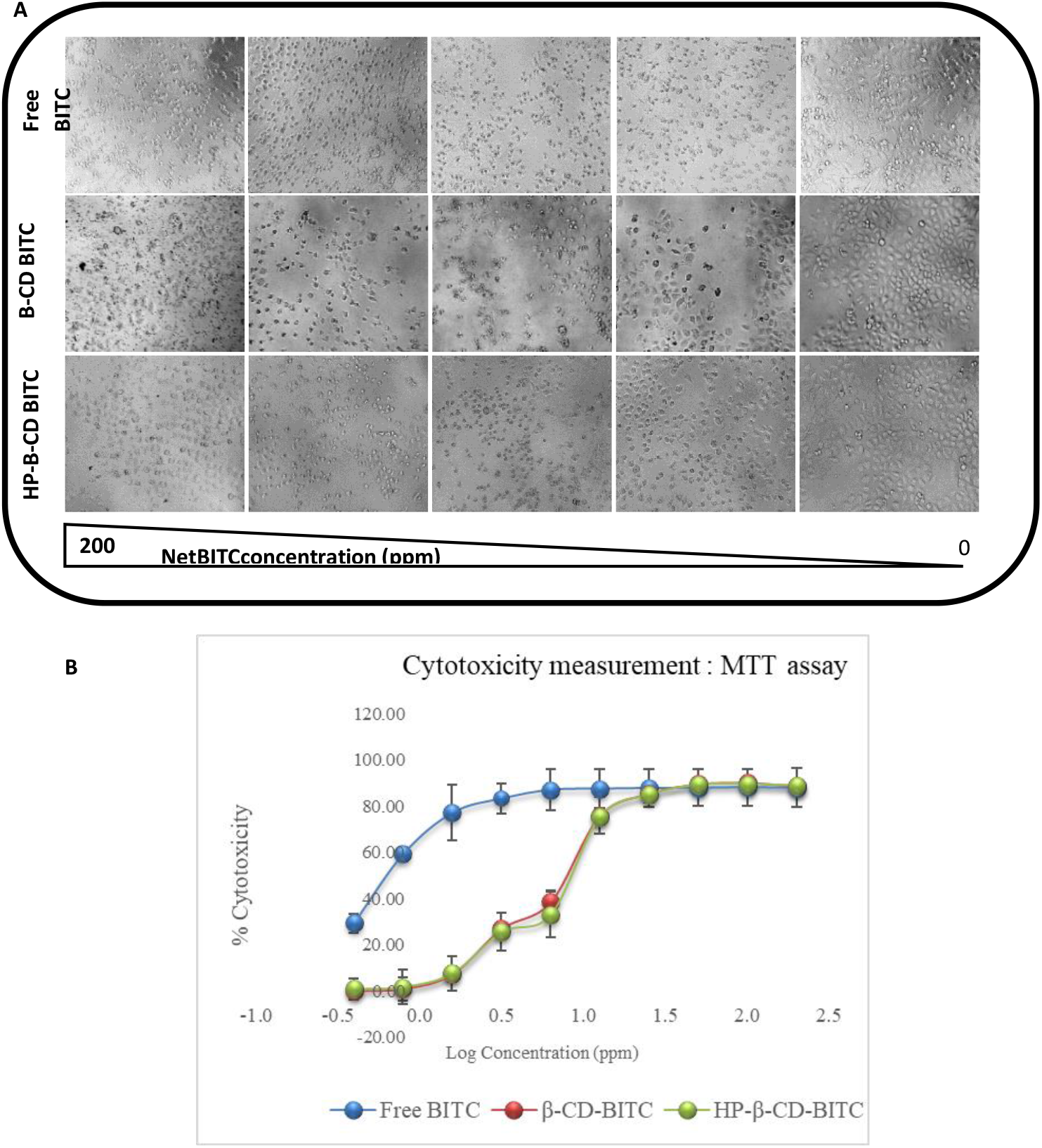
Cell viability of MDA-MB-231 cells. A) Representative photomicrographs showing the morphology of MDA-MB-231 cells treated with Free BITC, β-CD-BITC, and HP-β-CD BITC. Images were acquired 24 h post-treatment. Images reveal concentration-dependent toxicity in all the treatments. Image magnification 10X. Top row: Cells treated with free BITC, Middle row: β-CD-BITC, Bottom row: HP-β-CD-BITC. B) Cell cytotoxicity vs. concentration scatter plot. BITC showed maximum toxicity. β-CD-BITC and HP-β-CD-BITC showed similar toxicity. IC50 values for all the treatments are shown on the right-hand bottom of the plot. The data shown are the mean of 3 independent experiments, and error bars indicate standard deviation.

Figure 4 (B) depicts the annihilation of the cell (i.e., cytotoxicity) following their exposure to different concentrations of BITC and the fabricated complexes. Free BITC was found to be highly cytotoxic as compared to both β-CD-BITC and HP-β-CD BITC, and respective mean IC_50_ values (SD) were 0.6324 (0.44), 7.33 (1.68) and 7.887 (2.17) ppm. Dose-dependent cell viability was observed for all the test samples. β-CD-BITC and HP-β-CD-BITC showed similar toxicity. Both the formulations were able to show the sufficient levels of cell death in MTT based assessment of cell viability; however, it was apparently low as compared to bare form even when these were leveled to have equivalent concentrations. This being obvious led us to use both the formulation at higher concentrations in subsequent experiments. Morphology images show rounded floating cells at higher concentration treatments, and the number of such events was decreased with a fall in the equivalent concentration of BITC. Images were acquired 24 h post-treatment. Images revealed concentration-dependent toxicity in all the treatments. The data shown are the mean of three independent experiments, and the error bars indicate standard deviation. Based on the toxicity response of all the treatments, the rest of the experiments were performed at 2.5, 5, 10, and 20 ppm equivalent concentrations of BITC.

### Cellular uptake

Intracellular localization of free BITC, β-CD-BITC, and HP-β-CD BITC was evaluated using confocal microscopy and flow cytometry. Fluorescence, although not very bright in intensity, being another important property, acted instrumental in tracking its intracellular presence without any further fluorochrome stapling and was used to detect the cellular uptake. This has been depicted in Fig. 5. The left column in Fig. 5 (A) presents the Differential interference contrast images (DIC), followed by BITC fluorescence in 530 nm in the middle column, and lastly, the last column shows an overlay of DIC and fluorescence. Native BITC and both inclusion complexes made from β- and HP-β-cyclodextrins displayed green fluorescent emission in the cytoplasmic region of the cells. The cell nucleus was stained with DAPI to differentiate and locate any nuclear localization. However, it could not detect at least using confocal microscopy. The effectively augmented cellular uptake of BITC can be ascribed to the nano-CD carriers, which facilitate the drug transportation across the cell membrane (40).

**Figure 5:**
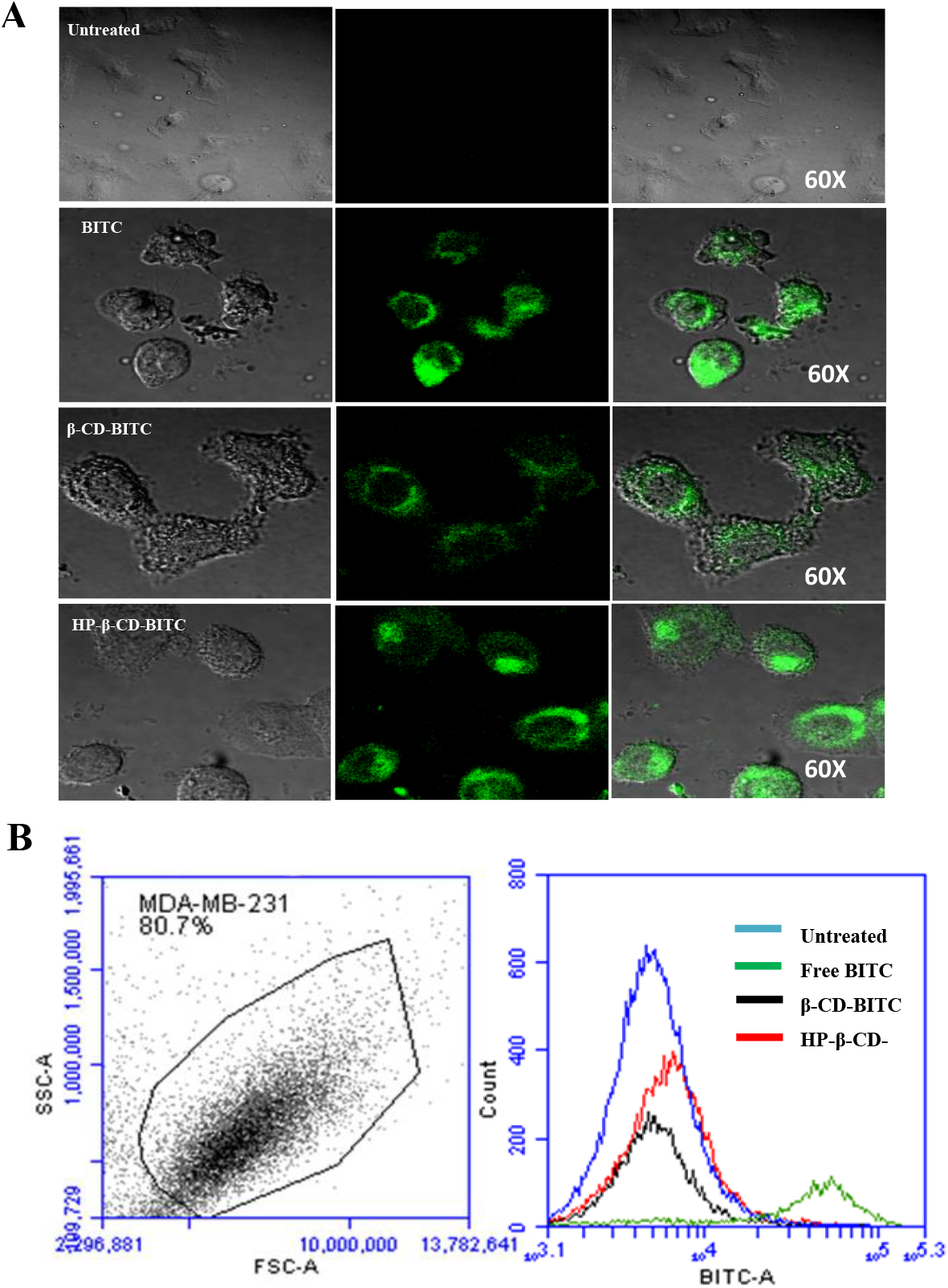
Cellular uptake of BITC in treated MDA-MB-231 cells. A) Confocal images of control cells, cells treated with Free BITC, β-CD-BITC and HP-β-CD-BITC. Left column in A) are DIC images, Middle column, BITC fluorescence in 530 nm channel and Right column shows overlay of DIC and fluorescence. B) Quantitative analysis of cellular uptake using flow cytometry. Left plot in B) shows dot plot for selection of MDA-MB-231 cells by creating gate and right plot in B) shows overlay of histogram from cells individually treated with Free BITC, β-CD-BITC and HP-β-CD-BITC or control cells

Flow cytometry analysis (Fig. 5 (B)) was performed to quantify cellular uptake. The results exhibited that free BITC treated cells had maximum uptake, followed by both inclusion complex and not significantly differing from each other. This quantized data on cellular uptake is clearly in line with the cell viability study and supports the higher half maximal inhibitory concentration IC_50_ values observed for free BITC. Higher accumulation of BITC as compared to both β-CD-BITC and HP-β-CD BITC, suggest the lower IC_50_ observed for BITC indirect administration. BITC given directly can cause massive cell death-like havoc, and it was evident in apoptosis assay; however, this extreme behavior was concentration-dependent, and when BITC was encapsulated in the cyclodextrins, it caused a restriction on its release in the given time and space.

### Phosphatidylserine (PES) externalization and apoptosis for cell death mechanism

Annexin-V binds to PES in calcium on the outer cell membrane, which otherwise present at the inner leaflets (41). This externalization, which indicates apoptosis, was detected using fluorescently labeled Annexin-V staining (Fig. 6). This is particularly vital as cellular metabolism gets affected by the drugs, but the plasma membrane remained relatively undamaged. In the absence of phagocytes, cells that show late apoptotic events, also known as necroptosis, were identified by including propidium iodide, which labels such cells due to the compromised cell membrane. MDA-MB-231 breast cancer cells were exposed to BITC, and a stoichiometrically equivalent amount of BITC in complexes (β-CD-BITC and HP-β-CD BITC) and apoptosis were quantified employing flow cytometry. The probability of apoptosis caused by BITC, β-CD-BITC, and HP-β-CD BITC, especially at their lower concentrations, was estimated qualitatively by flow cytometry, and all treatments showed a dose-dependent increase in apoptotic population. Along with concentration-dependent behavior, it was also noteworthy that cells were dying through the apoptotic cell death mechanism as a necrotic population that can cause significant immune reactions was merely observed. MDA-MB-231 cells, like any cancer cells, have continuous proliferation property acquired through continuous cell cycling behavior. Apoptosis induced by BITC in these cells could be attributed to a block in the cell cycling and was perceived at the G2/M phase of the cell cycle. Although free BITC leads the board and almost 90% of cells were apoptotic at 10 ppm, both β-CD-BITC and HP-β-CD BITC also showed concentration-dependent apoptotic events (Fig. 6). It was worth noting that concentration above 10 ppm also increased the necrotic population in all the treatments, which could be due to the dose-dependent lethality of BITC.

**Figure 6:**
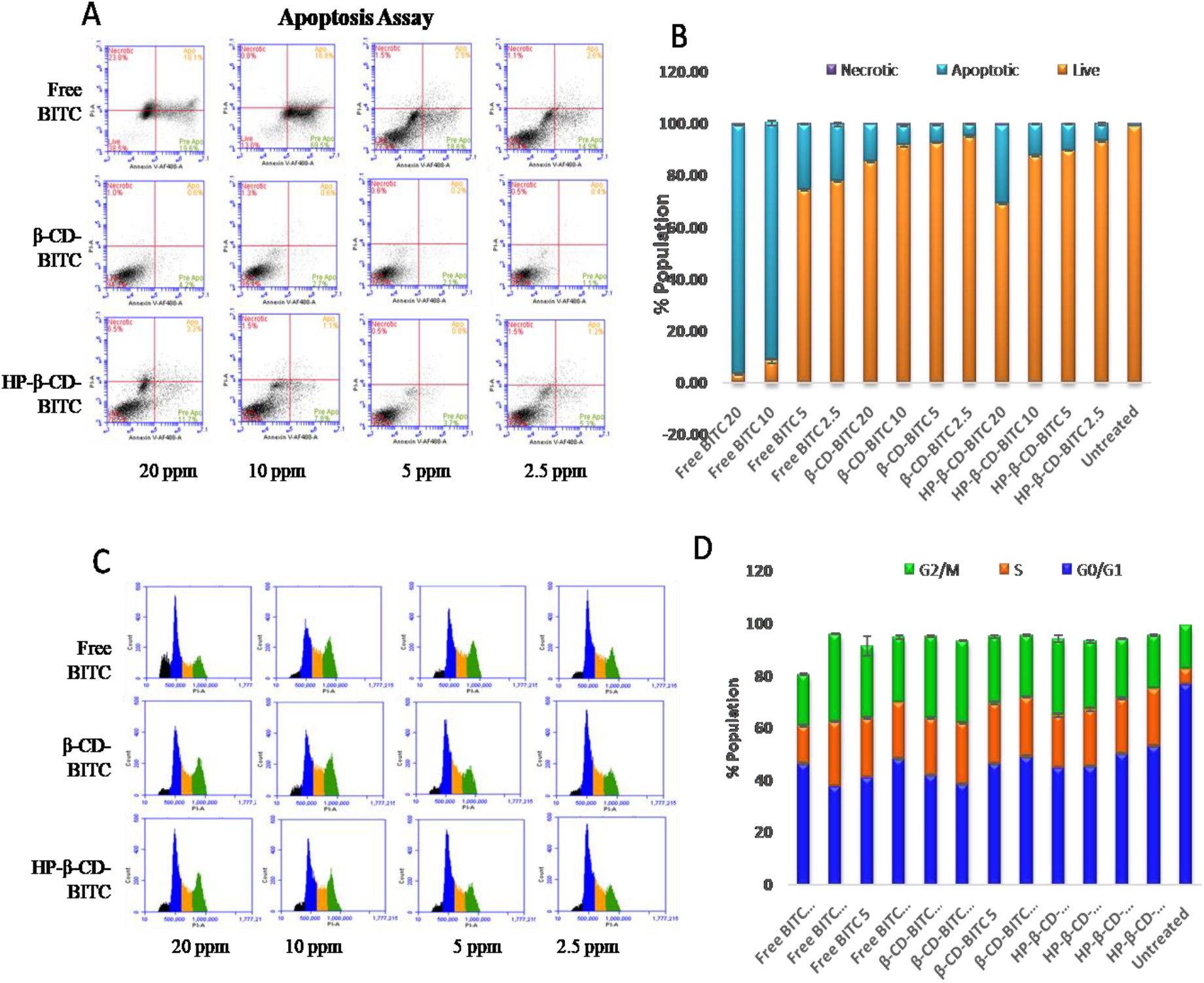
Cell death mechanism analysis in MDA-MB-231 cells. A) Flow cytometry dot plots showing Annexin-Propidium iodide uptake analysis for measurement of apoptotic and necrotic cell population after 24 hrs of respectively mentioned treatment in MDA-MB-231 cells. B) Bar graph showing quantitation of percentage apoptotic and necrotic cell population in the respective treatment group; Cell cycle analysis in MDA-MB-231 cells. C) Flow cytometry histogram plots showing the distribution of various cell cycle phases in the MDA-MB-231 cell population. In A), Blue color shows cells in G0/G1 phase, yellow color shows S-Phase, green indicates G2/M, and black indicates Sub-G0/G1 fraction, which indicates dead population. D) Quantitative display of various cell cycle phases in respectively treated cells. The result shown here indicates mean values along with the bar showing standard deviation.

### Cell cycle analysis

Malignant transformation is known to enhance cell cycling behavior in affected cells results in continuous proliferation. Many anti-cancer therapies act on one or more phases of the cell cycle, which results in either cytostatic or cytotoxic cellular events. Block in cell cycle phase can be observed as an increase in the number of cells in one of the specific cell cycle phase [37]. DNA content varies between these phases and can be used to identify different phases. Cells in G0/G1 phase have diploid (2n) DNA content, and once they cross the checkpoint, cells enter into the S phase and result in an increase of total DNA content and get doubled (4n) at the end of the S Phase. After complete duplication of DNA, cells enter into the mitotic phase. Propidium iodide, if facilitated into the cells by permeabilizing the cell membrane, can binds to DNA in a stoichiometric manner and can be analyzed on a flow cytometer. All the treatments showed concentration-dependent G2/M block in treated cells (Fig. 6). A similar arrest in the G2/M phase by BITC has been observed by Zhang *et al*. (42) in human pancreatic cancer cells. The maximum G2/M population was observed with a maximum concentration of treatment, i.e., 20 ppm. However, free BITC showed maximum G2/M arrest at 10 ppm, and a further increase in concentration resulted in the dead hypodiploid population present before the G0/G1 population. Arrested cells are known to dye because of the initiation of apoptotic events. If followed, cells appear to become hypodiploid and can be visualized to appear as a separate peak before G0/G1 in DNA histogram (Fig. 6).

### Reactive Oxygen Species (ROS) measurement

Cell’s mitochondria are the hub where ATP is synthesized by reducing molecular oxygen to water through electron transfer reactions. If not reduced completely, this results in superoxide anion radicals and other oxygen-containing radicals collectively known as ROS (43). Overproduction of ROS is deleterious to cell health and results in DNA-strand breaks, inflammatory responses and which finally leads to cell death. In the recent past, it has been shown that BITC can cause an increase in ROS after treatment, which could be responsible for the cell death in treated cells. Lin *et al*. (44)demonstrated the induction of ROS by BITC, which was found to be responsible for the apoptosis and autophagy in human prostate cancer cells. To evaluate the role of ROS in BITC induced cytotoxicity, monitored the levels of ROS using DCFHDA. BITC treatment induced significantly increased ROS in treated cells as compared to the basal level of ROS present, which was measured in untreated cells. Free BITC showed a continuous increase in ROS till 10 ppm, while both β-CD-BITC and HP-β-CD BITC showed low but modest ROS generation in a concentration-independent manner. It was also found that there is a concentration-dependent increase in ROS inside the cell. Strikingly it was observed that 20 ppm free BITC concentration showed less ROS, which could be due to the high cell death present in the condition, and probably most of the cells started dying even at the low time point (12hr) selected for the measurement of ROS. However, time could not be further reduced as it would appear as no visible effect in the rest of the treatment conditions. The results obtained are demonstrated in Fig.7. Increased ROS can cause the accumulation of lysosomes, which are hydrolases containing acidic organelles and increase in number and can apoptosis upon release of the contents in intracellular spaces.

### Lysosomes tracking and quantitation

Lysosomes are acidic vacuoles known to contain a variety of hydrolases and, when located intracellularly, can trigger apoptosis by releasing the content. Also, it is known that oxidative stress can induce lysosome-mediated cell death by destabilizing the lysosomal membrane (45). The permeabilization in the lysosomal membrane results in the release of internal contents and enzymes in the cytoplasm, which can be associated with cell death. To study the effect of BITC and BITC containing inclusion complexes on lysosomal enrichment, cells were stained with Acridine Orange (Himedia, India) for microscopic visualization or Lysotracker Red (Thermo Fisher, USA) for quantitation using flow cytometry. As the concentration of BITC increased, the lysosomal number was found to be increased in microscopic visualization (Fig. 7E). Also, flow cytometry quantization illustrated in Fig. 7(C) corroborated the same, and free BITC showed maximum lysotracker localization followed by β-CD-BITC and HP-β-CD BITC. Lysosomal targeting can be useful for especially apoptotic resistant cancer cells, and BITC induced lysosomal increase can add unique benefit to any conventional chemotherapy.

**Figure 7:**
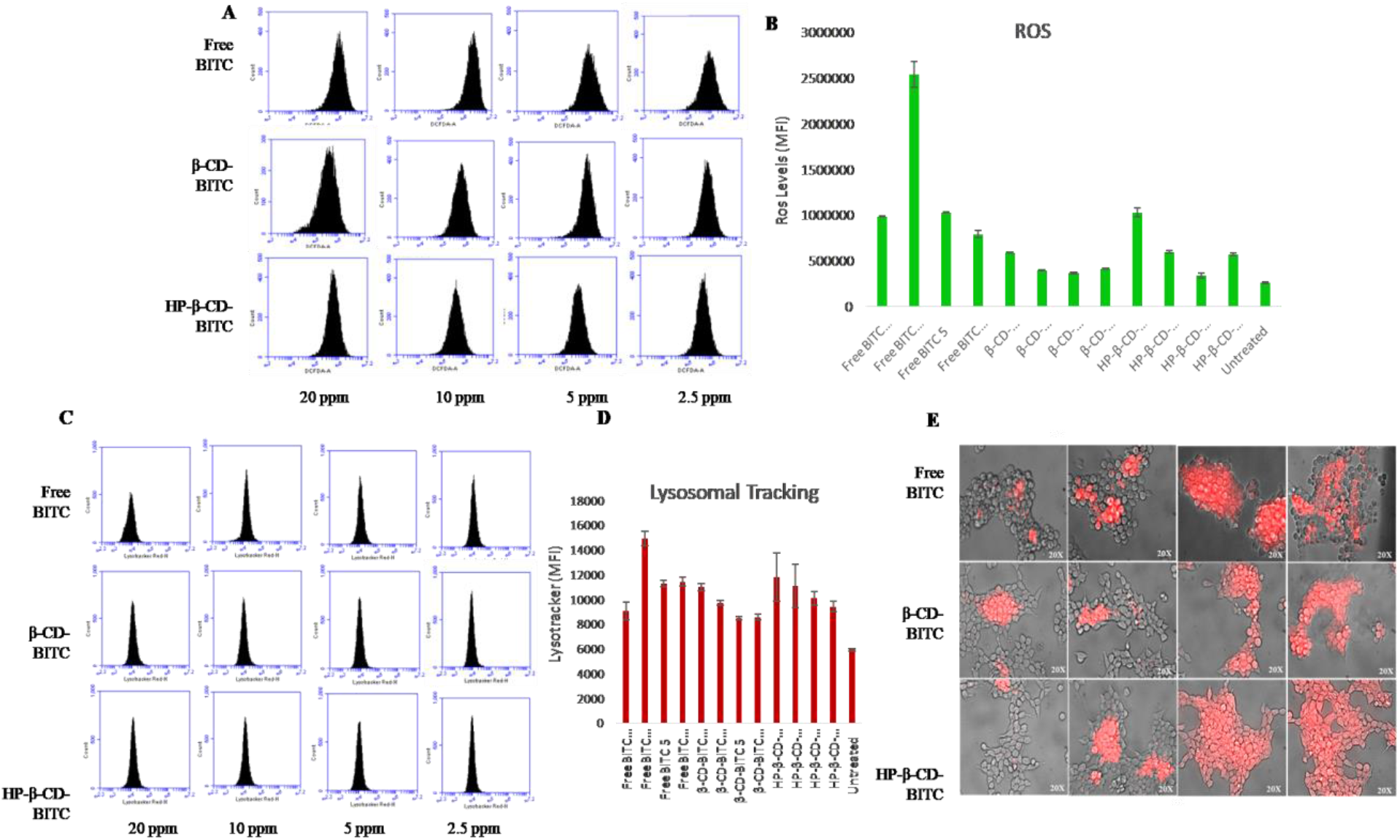
Reactive oxygen species production measurement in MDA-MD-231 cells. A) Flowcytometry histogram plots showing fluorescent intensity of DCFDA after different treatment in MDA-MD-231 cell. Right shift on x-axis indicates increase in fluorescence intensity which is directly proportional to presence of reactive oxygen species. B) Quantitative display of levels of ROS produced in respectively treated cells. Result show here indicates mean values along with bar showing standard deviation; Analysis of acidic vacuoles by lysotracker based flowcytometric analysis. in MDA-MD-231 cells. C) Flowcytometry histogram plots showing fluorescent intensity of lysotracker-red after various treatment in MDA-MD-231 cell. Right shift on x-axis indicates increase in fluorescence intensity of lysotracker red which is directly proportional to presence of acidic vacuoles. D) Quantitative display of levels of acidic vacuoles produced in respectively treated cells. Result show here indicates mean values along with bar showing standard deviation E) Acridine orange stained cells shows concentration dependent increase in acidic vacuoles (red) in respective treatment conditions.

### Wound healing scratch Assay

Continuous cell proliferation along with enhanced motility are two important hallmarks of aggressive cancer. BITC as an active constituent was shown to increase cell death and also expected to act as cytostatic at low concentration (section 4.4). Effect on cell migration using simple but powerful wound healing assay was assessed. Cell migration can be considered as an indicator for metastatic properties, and wound filling assay can be served as a surrogate to measure the potential of a drug compound (46). Purposefully free BITC was excluded from the analysis as BITC showed the higher toxicity in the apoptosis assay. It became pre-emptive that BITC will increase the open wound area because of higher toxicity and only, β-CD-BITC and HP-β-CD BITC were tested. There was a concentration-dependent interruption in an open wound infilling. The inhibition in the cell migration was observed clearly even after 24 h for all the tested concentrations for both the synthesized complexes. The obtained results promptly affirm the efficacy of the CDs in releasing BITC and attests the ability of the complexes to impede the growth of metastatic MDA MB 231 cells.

Furthermore, 20 ppm concentration showed an increase in the open wound space at a later time point due to induction of significant apoptosis. Results are presented as % open wound area against time and untreated cells, which shows maximum filling was used as a control (Fig. 8). Compared to the control, a significant decrease in the invading number of cells can be seen. Both β-CD-BITC and HP-β-CD BITC slowed the migration of cells, which could be either dependent or independent of the proliferation of these cells; however, current experimental conditions were not fully equipped to fully answer this query. Further studies are needed in this direction, although the obtained results substantiate the ability of the complexes to target the cell.

**Figure 8:**
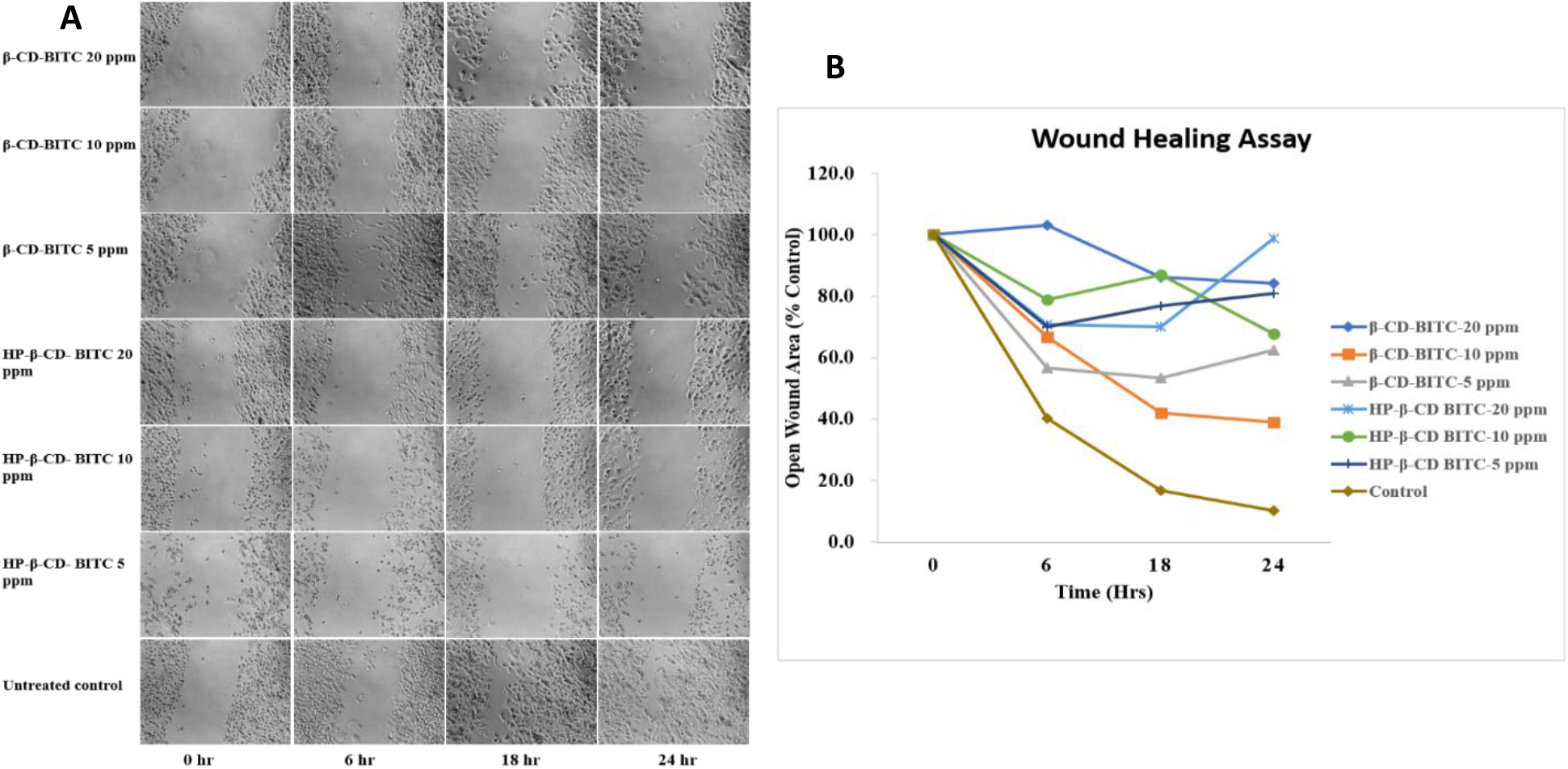
Analysis of cell migration properties in MDA-MB-231 cells in various treated cells group Bottom row in A) shows photomicrographs of untreated cells at 0, 6, 18 and 24 hours. Open gap which is called here as wound healing can be visualized in a time dependent manner. Other rows and columns show cells monolayer in variedly treated cells at different time points. B) % of open wound area which was calculated using T-scratch is plotted on a line plot to show comparative levels of cells migration in different groups.

## CONCLUSIONS

Nano-modules potentially provide a viable route to the potent yet fragile nutraceuticals. They can improve the effectiveness of the currently available cancer therapies by the conceptualization of chemoprevention using natural bioactive compounds. A successful attempt has been made to fabricate the sonochemical complexes of BITC with β-CD and HP-β-CD with effective antiproliferative activity under *in vitro* conditions. Aptly characterised employing various spectroscopic and microscopic methods, the complex between BITC and CDs (β-CD and HP-β-CD) was found to possess sustained release characteristics which would attribute significantly to improve the biopharmaceutic hurdles of BITC. Korsmeyer and Pappas equation in kinetic modeling showed that the transport mechanism of drug release of BITC from inclusion complexes is Fickian in nature. The *in vitro* assessments using MTT assay proved the antiproliferative effect of the BITC even in the complexed form.

Further, the cell cycle arrest studies and apoptosis assay elucidated the mechanism of cytotoxic potential of BITC. ROS generation studies and quantification of lysosomal tracking further provided details into the therapeutic activity of BITC. These results provide a promising platform for the development of more nano prototypes as efficient drug delivery systems for chemoprevention/therapy of BITC. Further work is necessary to testify the effectiveness of BITC in the nanocomplexes of CDs using *in vivo* models.

## ACKNOWLEDGMENTS

SKM expresses gratitude to CSIR for financial assistance in terms of a Major research project. KK would like to thank the DST for funding and INSPIRE faculty award. SU is thankful to CSIR for SRF (Open) fellowship.

