## Supplementary Information for "Synthesis, Physiochemical and Biological evaluation of Inclusion Complex of Benzyl Isothiocyanate encapsulated in cyclodextrins for triple negative breast cancer"

**Figure S1: DSC Thermograms of (A) BITC, β-CD BITC and β-CD (B) BITC, HP-β-CD BITC and HP-β-CD**

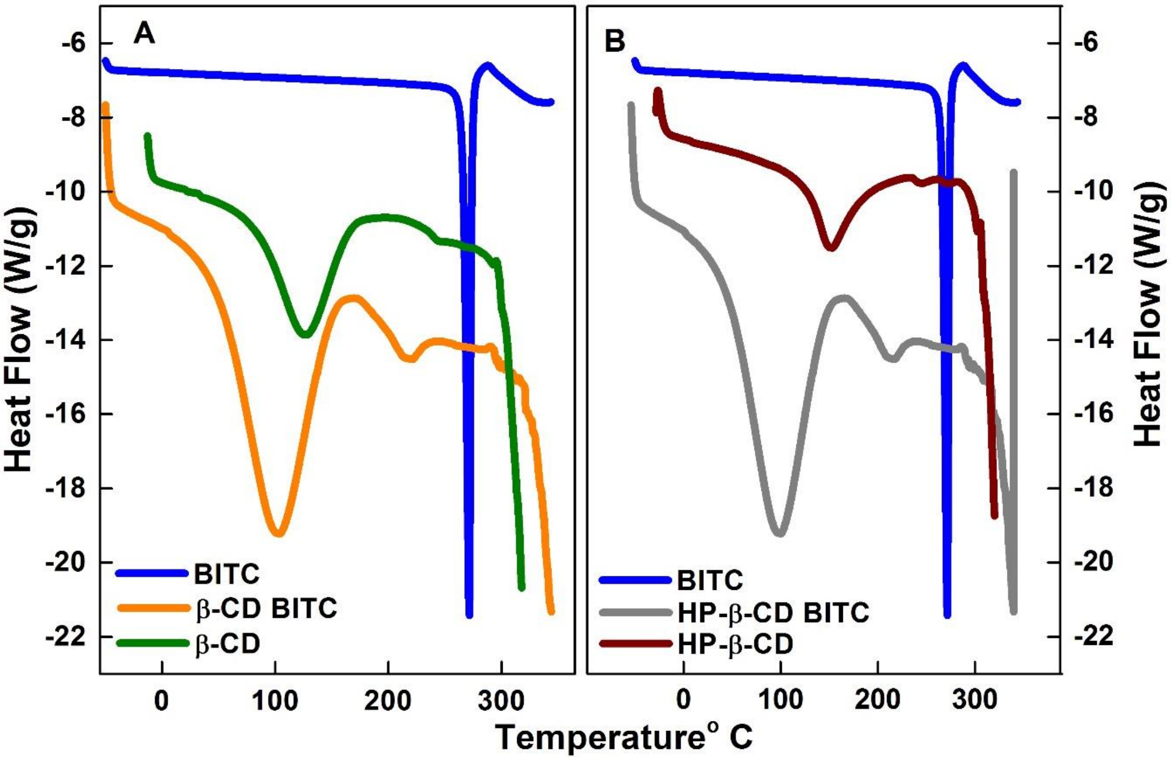

**Figure S2. Electron density profiles for β-CD BITC and HP-β-CD BITC**

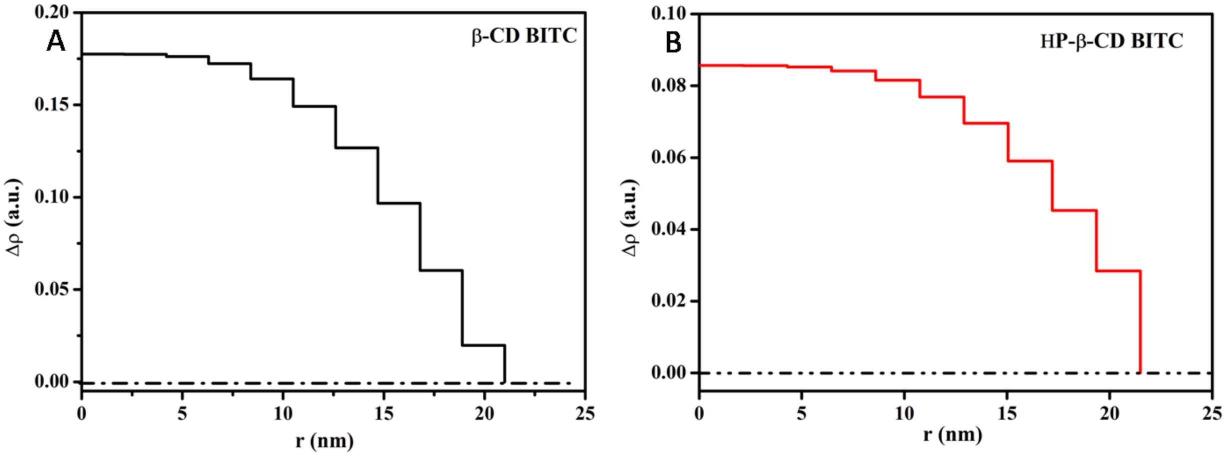

**Figure S3. Cumulative Release profile of BITC**

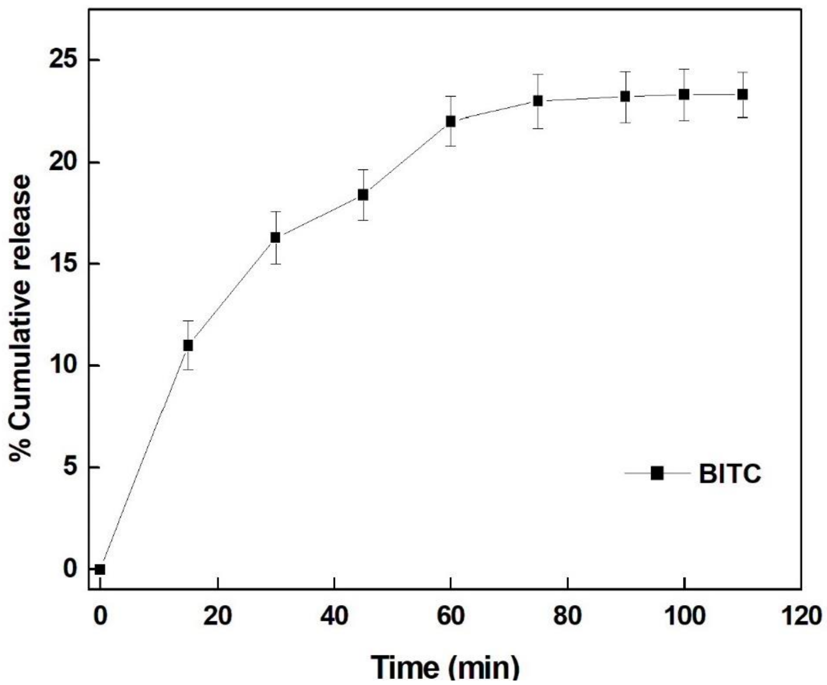

**Table S1 Kinetic Models for the release of BITC from β-CD and HP-β-CD at pH 7.2 and 5.5**

| Kinetic Model | | β-CD BITC (pH 5.5) | β-CD BITC (pH 7.2) | HP-β-CD BITC (pH 5.5) | HP-β-CD BITC (pH 7.2) |
|---|---|---|---|---|---|
| Zero order | $k_o$ | 0.235 | 0.202 | 0.275 | 0.249 |
| | $R^2$ | 0.681 | 0.756 | 0.654 | 0.556 |
| First order | $k_1$ | 0.004 | 0.003 | 0.006 | 0.005 |
| | $R^2$ | 0.895 | 0.903 | 0.925 | 0.853 |
| Higuchi | $k_H$ | 3.795 | 3.234 | 4.449 | 4.072 |
| | $R^2$ | 0.942 | 0.936 | 0.974 | 0.935 |
| **Korsmeyer-Peppas** | $\mathbf{k_P}$ | **4.265** | **4.591** | **5.996** | **6.294** |
| | **N** | **0.478** | **0.441** | **0.444** | **0.419** |
| | $\mathbf{R^2}$ | **0.992** | **0.993** | **0.994** | **0.989** |
| Hixson-Crowell | $k_{HC}$ | 0.001 | 0.001 | 0.002 | 0.001 |
| | $R^2$ | 0.843 | 0.873 | 0.870 | 0.780 |
